# Punctuational evolution is pervasive in distal site metastatic colonization

**DOI:** 10.1101/2024.04.08.588529

**Authors:** George Butler, Sarah R. Amend, Robert Axelrod, Chris Venditti, Kenneth J. Pienta

## Abstract

The evolution of metastasis represents a lethal stage of cancer progression. Yet, the evolutionary kinetics of metastatic disease remain unresolved. Here, using single cell CRISPR-Cas9 lineage tracing data, we show that in metastatic disease, gradual molecular evolution is punctuated by episodes of rapid evolutionary change associated with lineage divergence. By measuring punctuational effects across the metastatic cascade, we show that punctuational effects contribute more to the molecular diversity at distal site metastases compared to the paired primary tumor, suggesting qualitatively different modes of evolution may drive primary and metastatic tumor progression. This is the first empirical evidence for distinct patterns of molecular evolution at early and late stages of metastasis and demonstrates the complex interplay of cell intrinsic and extrinsic factors that shape lethal cancer.

---

Cancer is an evolutionary process ^1^ in which genetic and epigenetic alterations accumulate over time, driving molecular divergence and widespread genetic intratumor heterogeneity ^2^. As a result, multiple studies have sought to characterize the variation in the rate of evolution (the tempo) and the distribution of evolutionary rates through time (the mode) during primary tumor progression ^3–7^. Yet, the primary cause of cancer related mortality is due to the evolution of metastasis ^8^, the spread of cancer to different sites within the body. Despite the clinical importance, the evolutionary roadmap of metastatic disease remains poorly understood ^9^.

The spatial and temporal heterogeneity of metastatic disease creates both analytical and practical challenges for evolutionary characterization ^10^. However, recent efforts with single cell CRISPR-Cas9 lineage tracing technologies have helped overcome these issues allowing for metastatic progression to be investigated at an unprecedented resolution and scale ^11,12^. Yet, no study to date has measured the temporal diversity, the mode, in the evolution of metastatic disease at the single cell level. For example, it is not clear whether the change in anatomical location from the primary tumor to a metastatic site is associated with increased heterogeneity in the distribution of molecular evolutionary rates through time. That is, given the small initial population size and change in microenvironment at a distal site, we might expect lineage diversification to be associated with molecular change, a dynamic known as punctuational evolution ^13^.

Punctuational evolution was originally proposed to explain the discontinuity of phenotypes within the fossil record ^14^. Specifically, Eldredge and Gould coined the term *punctuated equilibria* in which evolutionary change is concentrated at the point of speciation followed by a period of evolutionary stasis ^14,15^. In turn, subsequent experimental studies in bacteria showed that punctuational patterns of evolution at the phenotypic level can also be mirrored at the molecular level ^16,17^. However, in contrast punctuational evolution at the phenotypic level, punctuational evolution at the molecular level does not require a period of statis, just an association between molecular divergence and lineage diversification ^13^. Multiple studies have since leveraged branch scaled phylogenetic trees to show that punctuational effects at the molecular level are widespread across different evolutionary systems, including linguistic ^18^, species ^13^, and viral evolution ^19^. Yet in cancer, punctuated-like patterns of evolution are typically considered as early events in tumor progression and are studied in the context of large-scale karyotypic alterations or copy number variations ^20,21^, thus potentially missing ongoing patterns of punctuational evolution driven by changes in the microenvironment as well as population size or structure. As a result, the presence punctuational evolution, defined as the association between lineage divergence and lineage diversification in cancer, and specifically metastasis, remains unknown.

Here, we use single cell lineage tracing data ^12^ in combination with a Bayesian phylogenetic framework ^13^ to test for evidence of punctuational evolution during metastatic progression. If punctuational effects are present, we expect to find a positive association between the amount of molecular divergence through time and the net increase in lineage divergence. In contrast, if punctuational effects are not present, we expect to find no association between the amount of molecular divergence through time and the net increase in lineage divergence (Figure. 1).

**Figure. 1.**
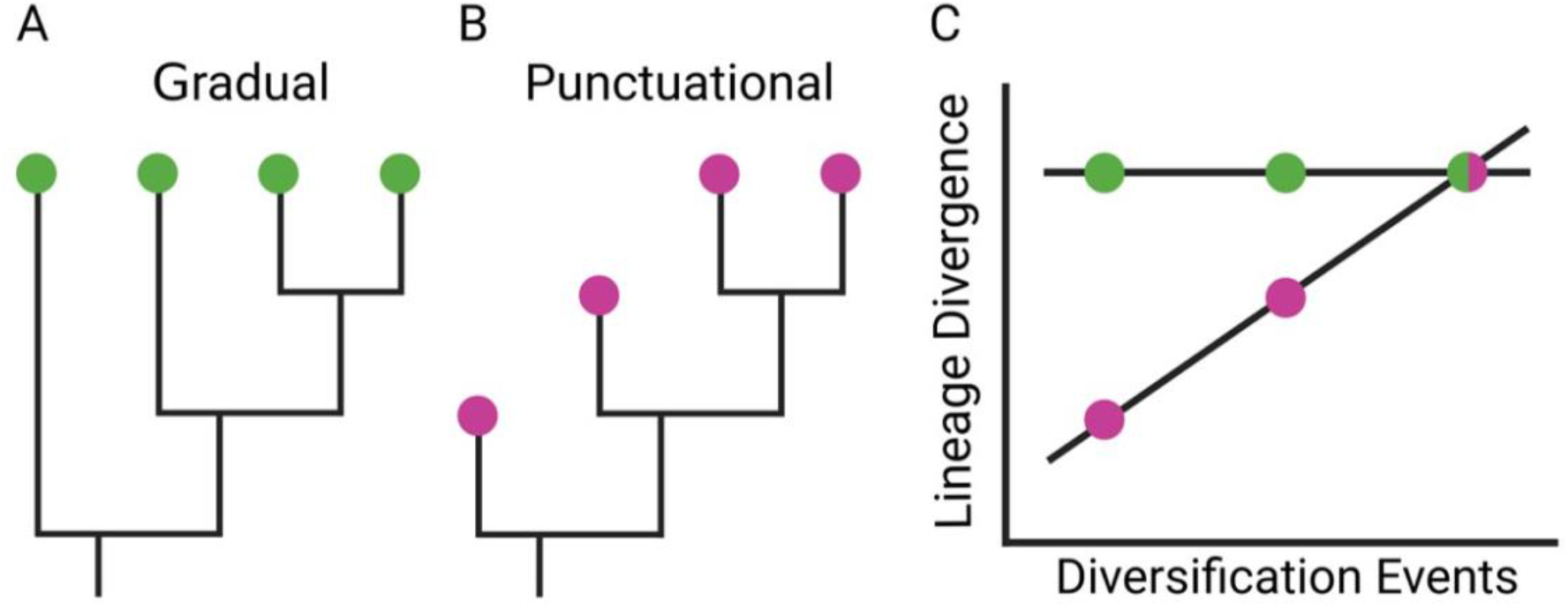
Detecting gradual and punctuational evolution. A) A gradual model of evolution assumes that molecular change, represented by branch lengths on the phylogenetic tree, is independent of lineage diversification. B) A punctuational model of evolution assumes that molecular change occurs at the point of lineage diversification. C) The relationship between lineage divergence, the sum of the branch lengths from root-to-tip, the number of diversification events along the path, can be used to test for evidence of punctuational evolution. A gradual model of evolution predicts no association between lineage divergence and diversification events. In contrast, a punctuational model predicts a positive association between lineage divergence and diversification events.

## Reconstructing branch scaled lineage trees at the single cell level

All single cell data used throughout is available from Yang et al ^12^. Briefly, sequencing data was collected from a lung adenocarcinoma genetically engineered mouse model (GEMM) in which embryonic stem cells were engineered with an evolving CRISPR-Cas9 dynamic lineage tracing system and inducible *Kras* and *Trp53* mutations, henceforth referred to as KP. Further data was also collected from two GEMMs that harbored an additional inducible LKB1 or APC mutation, henceforth referred to as KPL or KPA. Testing for punctuational effects across the three different mouse models controls for the potential effect of the individual GEMM system. The final dataset consisted of 10 mice, including: one KP mouse, seven KPL mice, and two KPA mice. Across the 10 mice, target site information was recovered for 25 separate lineages (10 metastatic and 15 non-metastatic), spanning 66 individual tumors and 14,882 cells (see Methods).

To reconstruct the phylogenetic relationship in each lineage, we used the edited target site information in combination with a mixture-model-based approach of phylogenetic reconstruction fitted in a Reversible Jump Markov Chain Monte Carlo (RJMCMC) framework ^22^. The branch lengths in the reconstructed trees correspond to the amount of expected evolutionary divergence between pairs of individual diversification events, measured in units of nucleotide substitutions (Figure. 2A). The mixture-model-based approach is a modification of existing likelihood-based methods of CRISPR lineage reconstruction ^23,24^ as it allows for qualitatively different models of sequence evolution ^22^ to be estimated across different cut sites. This modification is important to control for the variability in the CRISPR editing process and to ensure that accurate branch lengths are recovered. In turn, we fitted a non-reversible substitution model ^25^ to account for the “locked-in” nature of the CRISPR substitutions at a given target site ^26^. Finally, we derived a sample of 1000 trees from the posterior distribution of each lineage to account for the inherent topological uncertainty in the estimation process ^27^ (see Methods).

**Figure. 2.**
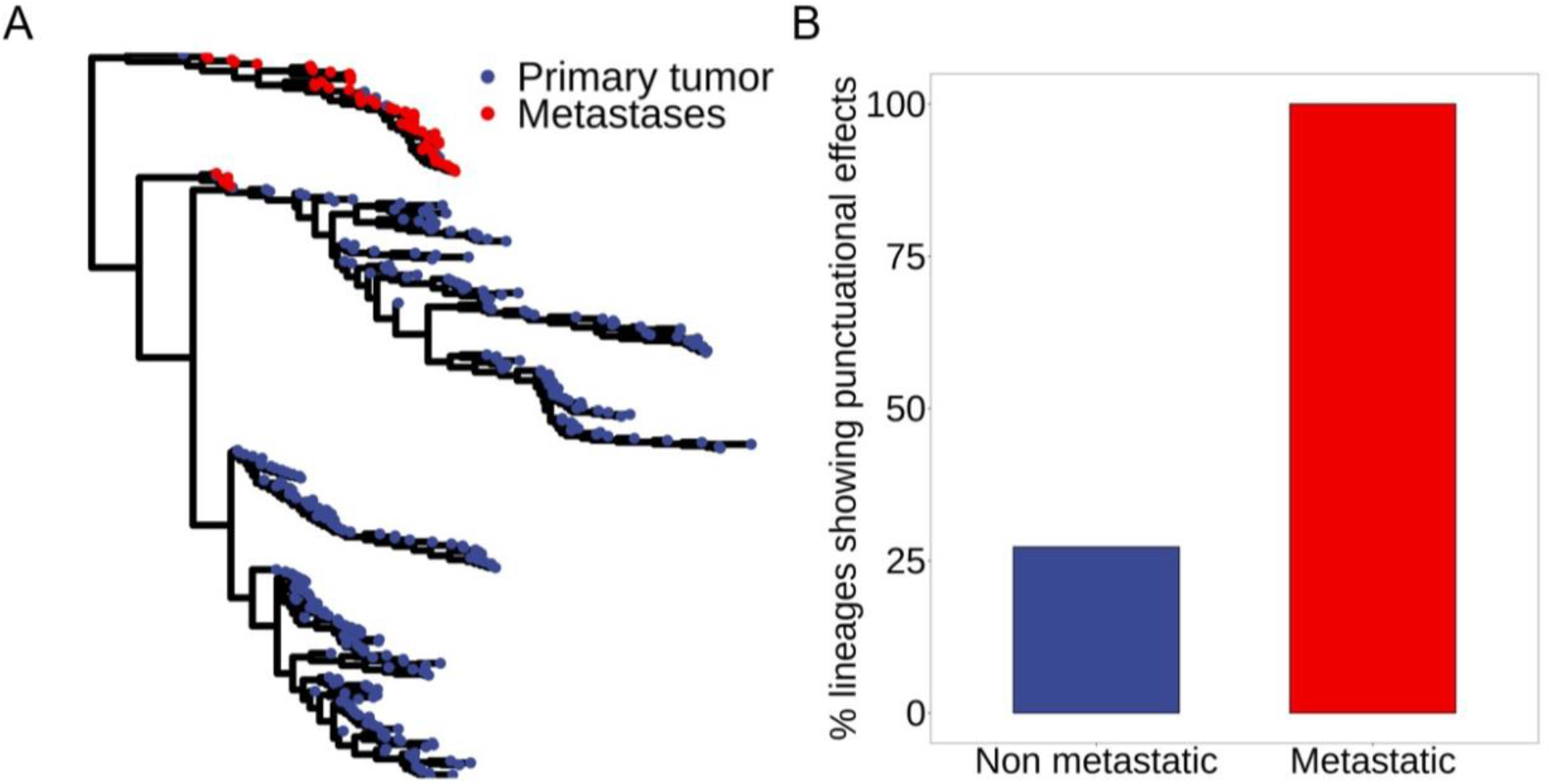
Punctuational evolution is more frequent in metastatic lineages. A) A metastatic single cell lineage tree. The blue and red points correspond to the individual cells recovered from the primary tumor and distal site metastases respectively. The branch lengths are proportional to the estimated kappa transformed (κ = 0.32) expected evolutionary divergence measured in units of nucleotide substitutions. B) The dataset consists of 18 lineages (7 metastatic and 11 non-metastatic). Punctuational evolution is significantly more common in metastatic compared to non-metastatic lineages (p = 0.011). All seven metastatic lineages show evidence of punctuational evolution but only three out of 11 non-metastatic lineages showed evidence of punctuational evolution (Table S1).

### Testing for evidence of punctuational evolution

To test for evidence of punctuational evolution, we tested for an association between the total lineage divergence and the total number of diversification events (one lineage splitting into two) ^28^. The total lineage divergence, LD, was calculated by summing the individual branch lengths from root-to-tip for each cell. Likewise, the number of diversification events, DE, was calculated by counting the number of nodes along the path from root-to-tip.

Next, we fitted a Phylogenetic Generalized Least Squares (PGLS) model in which lineage divergence was dependent on the number diversification events as well as the sum of the gradual effects in each branch along the path. That is, *LD =*β\**DE*+ *g* where β is the punctuational contribution to molecular divergence at each node and *g* is the gradual contribution. Under a gradual, homogenous model of evolution we expect to find no association between lineage divergence and diversification events and thus β = 0. In contrast, under a punctuational model of evolution in which diversification events are associated with molecular change, we expect to find a positive association between lineage divergence and diversification events and thus β > 0 (Figure. 1) ^13^.

We found a significant relationship between lineage divergence and diversification events, and thus evidence of punctuational evolution, in 17 of the 25 lineages (see Methods). However, seven of the 17 lineages with evidence of punctuational evolution also suffered from a well-known artifact of phylogenetic reconstruction, known as the node density artifact (see Methods, fig S1) ^29^, and were thus excluded from our analysis. As a result, our final dataset consisted of 18 lineages in which 10 of the lineages (7 metastatic and 3 non metastatic) showed evidence of punctuational evolution (Table S1). Finally, in the 7 metastatic lineages, we tested an alternative model in which a separate intercept and slope was estimated for metastatic cells within a given lineage. We found no significant difference in the estimated slopes and thus a single slope was estimated throughout (Table S2).

Evidence of punctuational evolution was present in both metastatic and non-metastatic lineages ranging in size from 247 to 1612 cells. However, we found that punctuational evolution was significantly more common in metastatic rather than non-metastatic lineages (p = 0.011, Figure. 2B). In fact, we found evidence of punctuational evolution in all seven metastatic lineages compared to only three of the 11 non-metastatic lineages. In contrast, while evidence of punctuational evolution was present in all three GEMMs (KP, KPL, and KPA), we found no association between the presence of punctuational evolution and the specific mouse model (p = 0.207, fig S2), suggesting that punctuational effects are independent of the given experimental mouse model.

Given that punctuational-like patterns of evolution in cancer are commonly considered with respect to karyotypic alterations and copy number variations ^20,21^, we fitted a secondary model in which the number of copy number variations (CNVs) was included as an additional independent variable (see Methods). We found no evidence of a significant association between lineage divergence and CNVs, yet the number of diversification events remained significant in the 10 lineages that had been identified previously (Table S3). Taken as a whole, these results suggest that punctuational effects are associated with metastatic progression and that punctuational patterns of evolution are not strictly a copy number-based phenomenon in cancer progression.

The slope between lineage divergence and the number diversification events (β) captures the increase in branch length (amount of molecular evolution) per diversification event ^13^. Yet, the absolute value of β will vary dependent on the lineage-specific overall rate of molecular evolution. That is, certain lineages may have a higher overall rate of molecular evolution compared to others. To allow for cross lineage comparisons, we used β to estimate the proportion of the total molecular diversity within each lineage that is attributed to punctuational effects (see Methods). Across the 10 lineages with evidence of punctuational evolution, the proportion of molecular diversity attributed to punctuational effects ranged from 0.011 - 0.125 with an average value of 0.06 (Table S1). That is, when punctuational evolution is present, an average of 6% of the molecular diversity is attributed to punctuational effects.

### Punctuational evolution has a greater impact on tumor progression at distal site metastases

Punctuational effects quantify the heterogeneity in evolutionary rates through time associated with lineage diversification. Two potential mechanisms of lineage diversification that may explain the presence of punctuational evolution are founder effects and ecological niche invasion ^13^.

Founder effects refer to instances in which a small subset of the population becomes spatially separated causing a temporary amplification in the effect of genetic drift prior to population expansion ^30^. In contrast, ecological niche invasion refers to adaptive evolution in which a subset of the population is better adapted to a set of new ecological stressors leading to a temporary increase in the rate of evolution at the point of lineage diversification ^31^. Crucially, both mechanisms are temporary ^32^, and both have the potential to act during cancer evolution, and specifically during the evolution of tumors at a distal site metastases. For instance, the number of cells that arrive at a distal site is expected to be smaller (generally thought to be a single cell or small cluster of cells) compared to the number of cells within the primary tumor (billions) ^33^.

Likewise, the environmental selective pressures at a distal site are expected to be different compared to the primary tumor microenvironment in which the cells first evolved ^34,35^. As a result, we hypothesize that punctuational evolution would have a greater impact on the evolutionary trajectory of tumor formation at a distal site compared to within the primary tumor. If true, we would expect to find a greater punctuational contribution to molecular diversity at a distal site compared to within the primary tumor.

To test whether the punctuational contribution to molecular diversity was different in the primary tumor compared to within distant site metastases, we partitioned the branches in each metastatic lineage according to whether the evolutionary change had occurred in the primary tumor or in a distal metastatic site. That is, each cell within a given lineages was labeled by location: primary tumor or distal metastatic site (e.g. lymph node, liver, bone, etc). In turn, for each branch in each tree in the posterior distribution for a given lineage, we compared the state of the ancestral and descendant nodes using a probabilistic method of ancestral state reconstruction to account for the uncertainty in the estimation process ^36^. If both the ancestral and descendant nodes were estimated to be in the same location, then the branch was categorized as that location (i.e, either primary tumor or metastatic). In contrast, if ancestral and descendant nodes were estimated to be in different locations, then the branch was categorized as a transition (Figure. 3A). Finally, we quantified the primary tumor and metastatic specific punctuational contribution within each lineage by calculating the ratio of primary tumor or metastatic branches with respect to the proportion of the total tree length attributed to either the primary tumor or distal site metastases (see Methods).

**Figure. 3.**
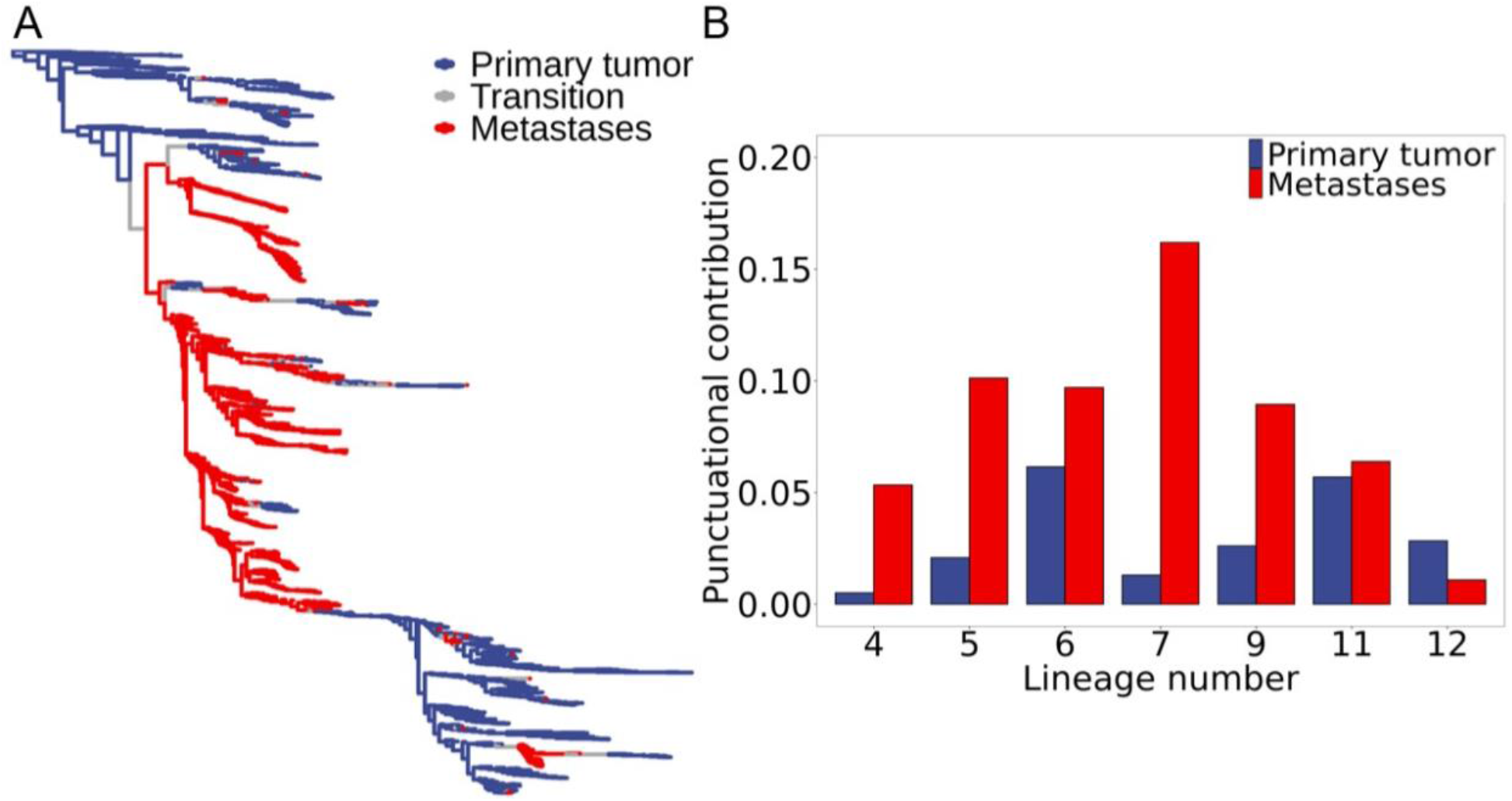
Punctuational effects contribute more to the molecular diversity in distal site metastases. A) A metastatic single cell lineage tree with inferred ancestral anatomical locations. The tree is transformed according to the estimated kappa value (κ = 0.31). The blue and red branches correspond to evolutionary change estimated to have occurred in the primary tumor and distal site metastases, respectively. The grey branches correspond to evolutionary change that has occurred in which the ancestral and descendent cells are estimated to be in different anatomical locations. B) The punctuational contribution to molecular diversity is significantly higher in distal site metastases (shown in red) compared to the paired primary tumor (shown in blue) in six of the seven metastatic lineages (Table S2).

We found that the punctuational contribution to molecular diversity was significantly higher in distal site metastases compared to the paired primary tumor in six of the seven metastatic lineages (Figure. 3B, Table S4). Moreover, we found that the proportion of molecular diversity attributed to punctuational effects ranged from 0.011 - 0.162 in distal site metastases compared to only 0.005 - 0.061 in the primary tumor. That is, 1 - 16% of the molecular diversity in distal site metastases is attributed to punctuational effects compared to only 0.5 - 6% of the diversity within the primary tumor. These results therefore suggest that punctuational evolution has a greater impact on the molecular landscape of distal site metastases compared to within the primary tumor.

Owing to the limited number of lineages with metastatic tumors of substantial size at different anatomical sites, we were unable to test whether punctuational effect sizes were differentially associated with specific metastatic sites. However, given that previous multi-region studies have shown that the lymph-node metastases emerge through wider evolutionary bottleneck compared to metastases at other distal sites ^33^, we would speculate that the punctuational contribution in lymph-node metastases would be smaller compared to other distal sites. Future investigation in this area is likely to result in further significant contributions to cancer biology. Nevertheless, these results here represent the first empirical evidence for qualitatively different modes of evolution at early and late stages of metastatic progression.

### Complex patterns of metastatic seeding are associated with stronger punctuational effects

Metastatic dissemination is a complex and inefficient process in which only a small fraction of the cells that leave the primary tumor ultimately seed and colonize at a distant site ^37,38^. A growing body of evidence suggests that patterns of metastatic seeding are considerably more complex than previously appreciated ^11,39,40^. Specifically, multiple studies have shown that a given metastatic site can be seeded by multiple independent lineages ^41^ and that cells from metastatic tumors can subsequently seed new metastases at different sites within the body ^42^. Given that punctuational effects cause deviations from gradual evolution ^13^, and our results show that punctuational effects have a greater impact on the trajectory of metastatic tumors, we speculate that more complex patterns of metastatic dissemination would result in stronger punctuational effects and thus greater deviations from gradual evolution.

To test the hypothesis that metastatic lineages seeded by a single monophyletic branch will have a smaller deviation from gradual evolution compared to lineages with polyphyletic seeding, we quantified the size of the deviation from gradual evolution by calculating the correlation coefficient, ρ, between lineage divergence and diversification (See Methods). We then characterized the level of “metastatic complexity” by estimating the degree of phylogenetic dispersion, D, ^43^ in the location of cells recovered from the primary tumor and distal site metastases. High metastatic complexity refers to instances in which metastatic cells are distributed throughout the tree (Figure. 4A and 3A). In contrast, low metastatic complexity refers to instances in which metastatic cells are clustered within a similar area(s) of the tree (Figure. 4B and 2A).

**Figure. 4.**
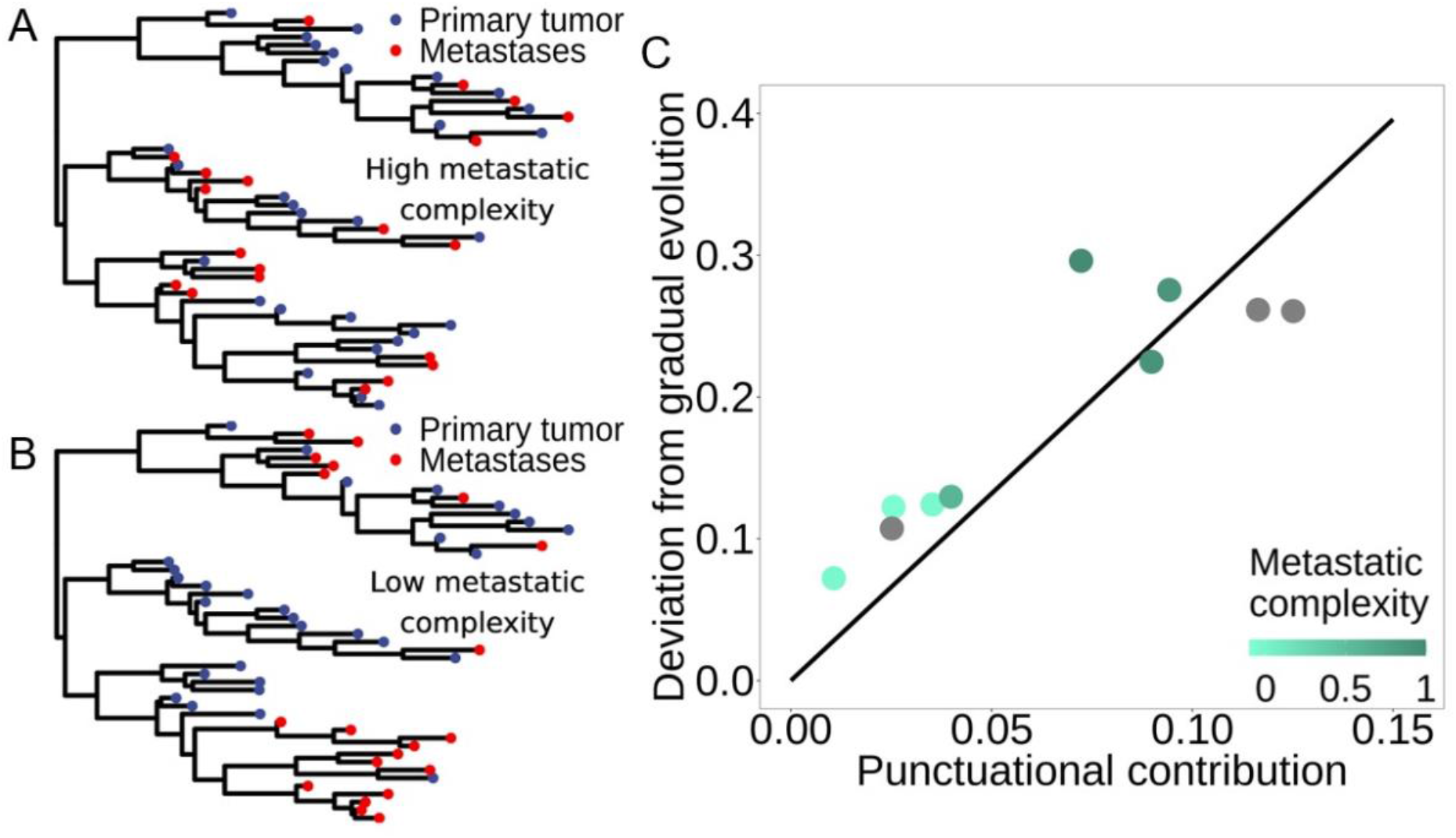
Larger deviations from gradual evolution are associated with stronger punctuational effects and increased metastatic complexity. A) A simulated tree with high metastatic complexity (D = 0.97) in which both the red and blue tips are distributed across the tree. B) A simulated tree with low metastatic complexity (D = -0.01) in which the red and blue tups are clustered within the tree. In both trees, the blue and red points correspond to simulated cells recovered from the primary tumor and distal site metastases. C) Metastatic lineages are shown in shades of turquoise and non-metastatic lineages are shown in grey. Darker shades of turquoise are associated with increased metastatic complexity, as measured by the degree of phylogenetic dispersion, D, in the location of cells within a given lineage that were recovered from the primary tumor and distal site metastases (see Methods). The fitted line shown in black is estimated from both metastatic and non-metastatic lineages and shows a significant positive association between the deviation from gradual evolution and the magnitude of the punctuational contribution (p < 0.001). Furthermore, lineages with increased metastatic complexity (shown in darker turquoise) are associated with larger deviations from gradual evolution.

First, we fitted a model across both metastatic and non-metastatic lineages in which deviations from gradual evolution are dependent on the size of the punctuational contribution. We found a significant positive association between the deviations from gradual evolution and punctuational contributions (p < 0.001, Figure. 4C). That is, stronger punctuational effects cause greater deviations from gradual evolution ^13^. Yet, along the punctuational continuum, we also found that greater deviations from molecular gradualism were present in lineages with increased metastatic complexity (Figure. 4C), suggesting that more complex patterns of metastatic seeding and reseeding are associated with increased heterogeneity in rates of molecular evolution through time.

## Conclusion

The evolution of metastasis signifies the evolution of lethal cancer. Yet despite the clinical importance, the evolution trajectory of metastatic disease remains broadly unknown ^9^. Here, we combined singe cell lineage tracing data ^12^ with robust phylogenetic comparative methods ^13^ to reveal that punctuational effects are a common molecular feature of metastatic progression.

Furthermore, by measuring punctuational effects throughout the disease course, we show that punctuational evolution has a greater impact on the evolutionary trajectory of distal site metastases compared to primary disease. Taken as a whole, these results represent the first empirical evidence of differential modes of evolution at early and late stages of metastatic progression and highlight the need to study metastatic dissemination as a continuous process rather that disjoint stages. Moreover, these results provide a tantalizing window into the breath of evolutionary dynamics that develop as a function of the interplay between a cancer cell and its environment ^35^, paving the way for future studies to investigate how exogenous factors such as systemic therapy differentially shape the trajectory of lethal disease progression.

## Materials and Methods

### Data and code availability

Raw single cell Target Site (for lineage reconstruction) and RNA-seq (for measuring copy number status) libraries as well as processed single cell clonality and sample information (verified from MULTI-seq and Lenti-Cre-BC libraries) are available from Yang et al *(12)*. Briefly, sequencing data was collected from a lung adenocarcinoma genetically engineered mouse model (GEMM) in which embryonic stem cells were engineered with an evolving CRISPR-Cas9 dynamic lineage tracing system and inducible *Kras* and *Trp53* mutations, henceforth referred to as KP. Further data was collected from two GEMMs that harbored additional inducible LKB1 or APC mutation, henceforth called KPL or KPA. Information regarding library construction, embryonic stem cell engineering, and sample preparation are detailed in Yang et al *(12)*.

All original code is available on GitHub at https://github.com/george-butler/punctuational_evolution_cancer. The software BayesPhylogenies *(22)* used for lineage reconstruction is available at https://www.evolution.reading.ac.uk/BayesPhy.html. The software BayesTraitsV4 *(44)* used to test for evidence of punctuational evolution and reconstruct ancestral states is available at https://www.evolution.reading.ac.uk/BayesTraitsV4.1.1/BayesTraitsV4.1.1.html.

### Single cell data preprocessing

#### Target site preprocessing

Target site libraries were processed using a modified version of the Cassiopeia *(45)* preprocessing pipeline (version 2.0.0) to retain the 296 base pair target sites in the final output table. The Cassiopeia parameters and threshold values were kept the same as outlined in Yang et al *(12)*. The preprocessed clonality and sample information (outlined above) was used to assign each of the target sites to a given lineage. The term “lineage” is used throughout to refer to individual cells that are descendant from the same original embryonic stem cell (as verified from the Lenti-Cre-BC library). To be included in the final dataset, a given lineage needed to have been recovered from a mouse that harbored at least one metastatic and one non-metastatic lineage. This criterion was used to try and control for between mouse differences that may affect the likelihood of metastatic progression. The final dataset consisted of 10 mice, including: 1 KP mouse, 7 KPL mice and 2 KPA mice. Across the 10 mice, 25 separate lineages were recovered (10 metastatic and 15 non-metastatic), spanning 66 individual tumors and 14,882 cells.

#### Target site alignment for phylogenetic reconstruction

Prior to phylogenetic reconstruction the target site information for each lineage was translated into a NEXUS file format. First, the 14-base-pair random integration barcode was removed from each target site, as well as the preceding 20 base pairs, resulting in a 262-base-pair sequence containing the three individual cuts. On average, 10 unique target sites were recovered for each lineage (19 out of 25 lineages had 10 unique target site) and all lineages had at least 7 unique target sites. However, due to practical limitations, the number of target sites that were recovered for an individual cell within a given lineage varied and may have been lower than the maximum number of target sites within the lineage. For instance, 10 unique targets were recovered for a given lineage but only 7 of the 10 target sites were recovered for a specific cell within the lineage. As a result, the missing target sites for an individual cell were marked with a blank sequence to preserve indexing across the alignments. Finally, the target sites for each cell within a given lineage were concatenated into a single N*262 sequence where N is equal to the number of unique target sites in the lineage and the integration barcodes were used to preserve the ordering across the lineage.

#### Measuring copy number variations

Copy number variations (CNVs) were quantified as outlined in Yang et al *(12)*. Briefly, RNA-seq libraries were processed using the 10x CellRanger pipeline (version 2.1.1) with the mm10 genome build. CNVs were then inferred from the processed RNA-seq libraries via the InferCNV R package (version 1.10.1) in which the parameters and threshold values were kept the same as outlined in Yang et al. All tumors containing a minimum of 5 cells were processed independently with normal lung cells from each background (KP, KPL, KPA) used as a reference. The number of CNVs per cell was calculated by summing the number of predicted CNV regions. In total, the CNV status was quantified for 14,876 out of 14,882 cells for which Target site information was present.

### Lineage reconstruction

Lineage reconstruction is an analytically challenging process due to the complex evolutionary dynamics, such as homoplasy, that can emerge overtime within a population. However, CRISPR-Cas9 lineage tracing data also has a unique set of characteristics that further complicate the reconstruction process. For instance, once edited, substitutions are considered to be “locked in” at a given cut site. Second, substitution rates can vary greatly between different target sites, as well as between individual cuts sites within a given target site. Finally, large deletions can cause adjacent cut sites to be overwritten thus erasing previous substitutions *(26,46-47)*. Taken together, these characteristics create a heterogenous set of dynamics that need to be properly accounted for during the reconstruction process to ensure that accurate topologies and branch length estimates are recovered.

#### Capturing complex patterns of molecular evolution in lineage tracing data

To detect and account for the heterogeneous patterns of evolution in CRISPR-Cas9 lineage tracing data we used a mixture-model based approach within a Reversible Jump Markov Chain Monte Carlo (RJMCMC) framework *(22)*.

First, given that the starting unedited target site sequence is known, we fitted a General Time Non-Reversible (GTNR) substitution model to estimate the frequency and rate of substitutions at sites within the target site. The GTNR model has an asymmetric 4x4 substitution rate matrix, **Q**, allowing for independent rates of substitution to be estimated in each direction. That is, a different substitution rate can be estimated from A to C compared to from C to A *(25)*. This detail is important as it allows for maximal flexibility and accounts for the “locked in” nature of substitutions at a given target site *(26)*. Next, to account for faster and slower rates of substitution at individual sites, we used a four-part discrete gamma rate model *(48)* to scale the individual elements of the substitution rate matrix, **Q**.

To account for qualitatively different patterns of molecular evolution across different areas of the alignment, such as between different target sites or cut sites, we used a mixture-model approach to fit *j* different evolution evolutionary models. That is, we estimated *j* independent **Q** matrices and then calculated the weighted sum to find the likelihood at each position across the sequence alignment. Thus, in theory, an individual model of evolution can be estimated for each cut site, or even different parts of each cut site in the alignment. Finally, to determine how many **Q** matrices need to be estimated within a given lineage, we used a RJMCMC framework. The RJMCMC framework removes the need for *a priori* specification and instead simultaneously estimates the elements of each **Q** matrix whilst also estimating the number of individual **Q** matrices. In turn, this reduces model complexity and removes the need for *post-hoc* model comparisons thus ensuring that reconstruction is tractable for lineage(s) spanning thousands for individual cells. Further details regarding the mixture-model-based approach and the RJMCMC framework are detailed in Pagel and Meade *(22)*.

#### Bayesian lineage reconstruction

Lineage reconstruction was performed using the software BayesPhylogenies *(22)*. The original unedited target site sequence was set as an outgroup to root each of the 25 lineages. Default priors were used throughout and the MCMC chains were sampled every 100,000^th^ iteration after visual convergence was achieved. A 1000 tree posterior distribution was produced for each lineage and the outgroup was removed prior to all subsequent stages of analysis. We found that the number of **Q** matrices estimated for each lineage scaled with lineage size and ranged between 1 and 24, highlighting the heterogenous mix of evolutionary patterns that emerge overtime and underlining the importance of estimating different models of evolution.

### Detecting punctuational evolution

To test for evidence of punctuational evolution, we tested for an association between total lineage divergence, *LD*, and the number of net diversification events, *DE*. Specifically, we used the equation *LD =*β*\*DE+ g* where β is the punctuational contribution of lineage diversification to molecular evolution at each node and g is the gradual contribution, as described in Pagel et al *(13)*. Under a gradual model of evolution, we expect to find no association between lineage divergence and diversification events. Thus, β = 0 and the equation simplifies to *LD* = *g* meaning that a gradual model of evolution is sufficient to explain the lineage divergence over time. In contrast, under a punctuational model of evolution, we expect to find a positive association between lineage divergence and diversification and thus β > 0 *(13)*.

#### Quantifying punctuational effects

Lineage divergence, *LD*, was calculated for each cell by summing the individual branch lengths from root-to-tip for each tree in the posterior distribution using the distRoot function in the adephylo R package *(49)*. Similarly, the distRoot function was also used to count the number of diversification events, *DE*, along the path from root-to-tip for each cell in each tree in the posterior distribution. A Phylogenetic Generalized Least Squares (PGLS) model was fitted in a maximum likelihood framework using the software BayesTraits *(44)* to estimate β separately for each tree in the posterior distribution. Phylogenetic signal (λ) was estimated during the model fitting process *(50)*. Default parameters were used throughout, and each tree was scaled to have a mean branch length equal to 0.1.

#### Node density artifact

The node density artifact is a systematic error of phylogenetic reconstruction that can cause branch lengths to be underestimated in areas of the tree with fewer taxa *(29)*. As a result, the node-density artifact can bias regression estimates and cause a relationship to appear between the amount of inferred total lineage divergence, *LD*, and the number of net diversification events, *DE*.

To check for the presence of the node density artifact, we used the δ test which predicts a curvilinear relationship between lineage divergence and diversification events when the artifact is present (fig S1) *(28)*. Specifically, the δ test fits a model of the form *LD =* β*\*DE*^*1/*δ^ *+ g* where *LD* is the lineage divergence, β is the punctuational contribution, *DE* is the number of diversification events, δ is the exponential diversification event scaler, and *g* is the gradual contribution. We used the software BayesTraits *(44)* to estimate β and δ simultaneously for each tree in the posterior distribution. Default parameters were used throughout, and each tree was also scaled to have a mean branch length equal to 0.1. The node density artifact is considered present when β is significantly bigger than 0 and δ is numerically greater than one *(13,18)*.

#### Criteria for identifying lineages with punctuational effects

We classified a lineage as having evidence of the node density artifact if β > 0 and δ > 1 (see *Node density artifact)* in at least 50% of the posterior distribution *(13)*. A total of 7 lineages (3 metastatic and 4 non metastatic) were found to have evidence of the node-density artifact and were thus removed from subsequent stages of analysis. In the remaining 18 lineages, we classified a lineage as having evidence of punctuational evolution if β > 0, δ ≤ 1 (see *Node density artifact)* in at least 50% of the posterior distribution and that the average value of δ across the posterior distribution was < 1. We found evidence of punctuational evolution in 10 out of the 18 lineages (7 metastatic and 3 non metastatic).

#### Assessing regression parameter significance

Regression parameter significance was assessed by the proportion of trees in the posterior distribution in which the parameter was significant at the 5% level. A parameter was classified as significant if in at least 50% of the trees in the posterior distribution the parameter was significant at the 5% level *(13)*.

### Quantifying the punctuational contribution to molecular diversity

The punctuational contribution to molecular diversity was quantified as outlined in Pagel et al *(13)*. Briefly, a bifurcating tree has 2(*c –* 1) branches, where *c* is equal to the number of individual cells in the lineage. If the overall length of the tree, *T*, is equal to the sum of the individual branch lengths measured in unit nucleotide substitutions. Then the ratio 2(*c* – 1)β/*T* measures the proportion of the tree length, the total molecular diversity, attributed to punctuational effects where β is the punctuational contribution of lineage diversification to molecular evolution at each node. Thus, if all lineage divergence is due to gradual effects, then β = 0 and the punctuational contribution, 2(*c*– 1)β/*T*, also equals zero. In contrast, if no gradual effects are present, 2(*c*– 1)β = *T* and the ratio 2(*c* – 1)β/*T =* 1.

### Quantifying the punctuational contribution across the metastatic cascade

To test whether the percent punctuational contribution to molecular diversity was different in the primary tumor compared to within distant site metastases, we partitioned the branches in each metastatic lineage according to whether the evolutionary change had occurred in the primary tumor or a distal metastatic site. That is, for each branch in each tree in the posterior distribution for a given lineage we compared the state of the ancestral and descendant nodes using ancestral state reconstruction (ASR) (see *Inferring ancestral states)*. Once finished, the branch lengths were then grouped into one of four categories: primary tumor, distant site metastases, transition, or undefined (see *Categorizing anatomical branch locations)*. Finally, we quantified the primary tumor and metastatic specific punctuational contribution within each lineage and then tested for a significant difference by comparing the pairwise difference between the two posterior distributions (see *Comparing punctuational contributions within a lineage)*.

#### Inferring ancestral states

To infer the state of each internal node within the posterior distribution of a given lineage we fitted a continuous time Markov model *(36)* using the location for which each cell in the lineage had been recovered, e.g. primary tumor or distal metastatic site (any tumor that is not the primary tumor e.g. lymph node, liver, bone etc). The model was fitted in a maximum likelihood framework using the software BayesTraits *(44)* to simultaneously estimate the transition rate from the primary tumor (P) to a distant metastatic site (M) (q_PM_), and the reverse (q_MP_), whilst also reconstructing the ancestral states for each node in the tree. We fixed the state of the root equal to the primary tumor state (P) but set no *a priori* restrictions on either of the transition rates (q_PM_ and q_MP_). The phylogenetic parameter kappa (κ) was estimated for each tree during the model fitting process to transform the individual branch lengths *(50)*. Default parameters were used throughout, and each tree was scaled to have a mean branch length equal to 0.1.

#### Categorizing anatomical branch locations

We categorized branches by comparing the ancestral and descendant nodal states resulting from the ancestral state reconstruction (ASR) procedure. The state of an internal node was set equal to the state with the highest probability. If both states had equal probability, P_ASR_= 0.5, then the nodal state was marked as “undefined”. If both the ancestral and descendant nodal states were the same (e.g. ancestor=P and descendant=P or ancestor=M and descendant=M) then the branch was also assigned to the same state. If both the ancestral and descendant nodal states were different (e.g. ancestor=P and descendant=M or ancestor=M and descendant=P) then the branch was assigned as a transition. Finally, if either the ancestral or descendant nodal state was “undefined” then the branch was assigned as undefined.

#### Comparing punctuational contributions within a lineage

We quantified the primary tumor and metastatic punctuational contribution within a given lineage by calculating the ratio of primary tumor or metastatic branches with respect to the proportion of the total tree length attributed to either the primary tumor or distal site metastases. The formula to calculate the punctuational contribution is outlined above (See *Quantifying the punctuational contribution to molecular diversity)*. To be included in the subsequent comparison, each tree needed to have at least 10 primary tumor branches and 10 metastatic branches, and the estimated punctuational contribution needed to be less than 1. The terminal branches were excluded due to their short length and thus potential to bias the average punctuational contribution. However, the same qualitative results are obtained when the terminal branches are included. Finally, we used a multiple comparison test with a Bonferroni correction to compare the punctuational contribution between the primary tumor and distal site metastases.

### Quantifying deviations from molecular gradualism

We quantified the size of the punctuational departure from gradual evolution by estimating the correlation, ρ, between lineage divergence and the number of diversification events as outlined in Pagel et al *(13)*. Specifically, ρ = (βσ^2^_LD_+ σ_g,LD_) / ((β^2^σ^2^_LD_ + σ^2^_g_)^½^ (σ^2^_LD_)^½^) where σ^2^_LD_ is equal to the variance in the number of lineage diversification events, σ^2^_g_ is equal to the variance in the gradual effects, and σ_g,LD_ is equal to the covariance between the gradual component and the number of lineage diversification events, and is assumed to be zero. If no punctuational effects are present, β = 0, we do not expect to find any deviations from a clock-like tempo of evolution and thus no correlation between lineage divergence and diversification. In contrast, if all molecular divergence is due to the punctuational effects, then we expect to find a perfect positive correlation between lineage divergence and diversification. That is, if the variance in the gradual effects, σ^2^_g_, is equal to zero then ρ = βσ^2^_n_ βσ^2^ _n_= 1.

We used a maximum likelihood framework to estimate ρ separately for each tree in the posterior distribution using the software BayesTraits *(44)*. Phylogenetic signal (λ) was estimated during the model fitting process *(50)*. Default parameters were used throughout, and each tree was scaled to have a mean branch length equal to 0.1. The average correlation was then calculated across the posterior distribution.

#### Estimating metastatic complexity

We characterized the level of “metastatic complexity” within a given metastatic lineage by estimating the degree of phylogenetic dispersion, D, *(43)* in the location of cells recovered from the primary tumor and distal site metastases. Metastatic lineages seeded by a single monophyletic branch with limited reseeding will have a lower degree of phylogenetic dispersion compared to lineages with polyphyletic seeding and continued reseeding. We estimated D separately for each tree in the posterior distribution using the phylo.d function in the Caper R package *(51)*. The average D was then calculated across the posterior distribution.

## Author contributions

Conceptualization: GB, SRA, RA, CV, KJP. Methodology: GB, CV, KJP. Investigation: GB. Formal analysis: GB and CV. Funding acquisition: GB, SRA, CV, KJP. Writing – original draft: GB. Writing – review & editing: GB, SRA, RA, CV, KJP.

## Data and materials availability

All data are available and referenced in the main text or the supplementary materials.

## Supplementary Materials

Figures S1 to S2

Tables S1 to S4

## References

1. Greaves, M. & Maley, C. C. Clonal evolution in cancer. Nature 481, 306 (2012).

2. Turajlic, S., Sottoriva, A., Graham, T. & Swanton, C. Resolving genetic heterogeneity in cancer. Nat Rev Genet 20, 404–416 (2019).

3. Sottoriva, A. et al. A Big Bang model of human colorectal tumor growth. Nat Genet 47, (2015).

4. Jamal-Hanjani, M. et al. Tracking the Evolution of Non–Small-Cell Lung Cancer. New England Journal of Medicine 376, 2109–2121 (2017).

5. Spain, L. et al. Late-stage metastatic melanoma emerges through a diversity of evolutionary pathways. Cancer Discov CD-22-1427 (2023).

6. Frankell, A. M. et al. The evolution of lung cancer and impact of subclonal selection in TRACERx. Nature 616, 525–533 (2023).

7. Househam, J. et al. Phenotypic plasticity and genetic control in colorectal cancer evolution. Nature 611, 744–753 (2022).

8. Rogiers, A., Lobon, I., Spain, L. & Turajlic, S. The genetic evolution of metastasis. Cancer Res 82, 1849– 1857 (2022).

9. Nguyen, B. et al. Genomic characterization of metastatic patterns from prospective clinical sequencing of 25,000 patients. Cell 185, 563-575.e11 (2022).

10. Gui, P. & Bivona, T. G. Evolution of metastasis: new tools and insights. Trends Cancer 8, 98–109 (2022).

11. Quinn, J. J. et al. Single-cell lineages reveal the rates, routes, and drivers of metastasis in cancer xenografts. Science (1979) 371, eabc1944 (2021).

12. Yang, D. et al. Lineage tracing reveals the phylodynamics, plasticity, and paths of tumor evolution. Cell (2022).

13. Pagel, M., Venditti, C. & Meade, A. Large punctuational contribution of speciation to evolutionary divergence at the molecular level. Science 314, 119–121 (2006).

14. Eldredge, N. & Gould, S. J. Punctuated Equilibria: An alternative to phyletic gradualism. in Models in Paleobiology (ed. Schopf, T. J. M.) 82–115 (Freeman Cooper, 1972).

15. Gould, S. J. & Eldredge, N. Punctuated Equilibria: The tempo and mode of evolution reconsidered. Paleobiology 3, 115–151 (1977).

16. Lenski, R. E. & Travisano, M. Dynamics of adaptation and diversification: a 10,000-generation experiment with bacterial populations. Proceedings of the National Academy of Sciences 91, 6808–6814 (1994).

17. Elena, S. F., Cooper, V. S. & Lenski, R. E. Punctuated evolution caused by selection of rare beneficial mutations. Science 272, 1802–1804 (1996).

18. Atkinson, Q. D., Meade, A., Venditti, C., Greenhill, S. J. & Pagel, M. Languages evolve in punctuational bursts. Science 319, 588 (2008).

19. Surya, K., Gardner, J. D. & Organ, C. L. Detecting punctuated evolution in SARS-CoV-2 over the first year of the pandemic. Frontiers in Virology 3, (2023).

20. Baca, S. C. et al. Punctuated evolution of prostate cancer genomes. Cell 153, (2013).

21. Gao, R. et al. Punctuated copy number evolution and clonal stasis in triple-negative breast cancer. Nat Genet 48, 1119–1130 (2016).

22. Pagel, M. & Meade, A. A phylogenetic mixture model for detecting pattern-heterogeneity in gene sequence or character-state data. Syst Biol 53, 571–581 (2004).

23. Seidel, S. & Stadler, T. TiDeTree: A Bayesian phylogenetic framework to estimate single-cell trees and population dynamic parameters from genetic lineage tracing data. Proceedings of the Royal Society B: Biological Sciences 289, 20221844 (2022).

24. Feng, J. et al. Estimation of cell lineage trees by maximum-likelihood phylogenetics. Ann Appl Stat 15, 343– 362 (2021).

25. Kozlov, A., Alves, J. M., Stamatakis, A. & Posada, D. CellPhy: accurate and fast probabilistic inference of single-cell phylogenies from scDNA-seq data. Genome Biol 23, 37 (2022).

26. McKenna, A. & Gagnon, J. A. Recording development with single cell dynamic lineage tracing. Development 146, dev169730 (2019).

27. Pagel, M. & Lutzoni, F. Accounting for phylogenetic uncertainty in comparative studies of evolution and adaptation. in Biological Evolution and Statistical Physics (eds. Lässig, M. & Valleriani, A.) 148–161 (Springer Berlin Heidelberg, Berlin, Heidelberg, 2002). doi:10.1007/3-540-45692-9_8.

28. Webster, A. J., Payne, R. J. H. & Pagel, M. Molecular phylogenies link rates of evolution and speciation. Science 301, 478 (2003).

29. Venditti, C., Meade, A. & Pagel, M. Detecting the node-density artifact in phylogeny reconstruction. Syst Biol 55, 637–643 (2006).

30. Carson, H. L. & Templeton, A. R. Genetic revolutions in relation to speciation phenomena: The founding of new populations. Annu Rev Ecol Syst 15, 97–131 (1984).

31. Rolland, J. et al. Conceptual and empirical bridges between micro- and macroevolution. Nat Ecol Evol 7, 1181–1193 (2023).

32. Venditti, C. & Pagel, M. Speciation as an active force in promoting genetic evolution. Trends Ecol Evol 25, 14–20 (2010).

33. Reiter, J. G. et al. Lymph node metastases develop through a wider evolutionary bottleneck than distant metastases. Nat Genet 52, 692–700 (2020).

34. Wu, Y. et al. Spatiotemporal immune landscape of colorectal cancer liver metastasis at single-cell level. Cancer Discov 12, 134–153 (2022).

35. Massagué, J. & Ganesh, K. Metastasis-initiating cells and ecosystems. Cancer Discov 11, 971–994 (2021).

36. Pagel, M., Meade, A. & Barker, D. Bayesian estimation of ancestral character states on phylogenies. Syst Biol 53, 673–684 (2004).

37. Lambert, A. W., Pattabiraman, D. R. & Weinberg, R. A. Emerging biological principles of metastasis. Cell 168, 670–691 (2017).

38. de Groot, A. E., Roy, S., Brown, J. S., Pienta, K. J. & Amend, S. R. Revisiting seed and soil: Examining the primary tumor and cancer cell foraging in metastasis. Molecular Cancer Research 15, 361–370 (2017).

39. McPherson, A. et al. Divergent modes of clonal spread and intraperitoneal mixing in high-grade serous ovarian cancer. Nat Genet 48, 758–767 (2016).

40. Brown, D. et al. Phylogenetic analysis of metastatic progression in breast cancer using somatic mutations and copy number aberrations. Nat Commun 8, 14944 (2017).

41. Al Bakir, M. et al. The evolution of non-small cell lung cancer metastases in TRACERx. Nature 616, 534– 542 (2023).

42. Gundem, G. et al. The evolutionary history of lethal metastatic prostate cancer. Nature 520, 353–357 (2015).

43. Fritz, S. A. & Purvis, A. Selectivity in mammalian extinction risk and threat types: a new measure of phylogenetic signal strength in binary traits. Conservation Biology 24, 1042–1051 (2010).

44. Meade, A. & Pagel, M. BayesTraits. Preprint at https://www.evolution.reading.ac.uk/BayesTraitsV4.1.1/BayesTraitsV4.1.1.html (2023).

45. Jones, M. G. et al. Inference of single-cell phylogenies from lineage tracing data using Cassiopeia. Genome Biol 21, 92 (2020).

46. McKenna, A. et al. Whole-organism lineage tracing by combinatorial and cumulative genome editing. Science (1979) 353, aaf7907 (2016).

47. Jones, M. G., Yang, D. & Weissman, J. S. New Tools for Lineage Tracing in Cancer In Vivo. Annu Rev Cancer Biol (2023).

48. Yang, Z. Maximum likelihood phylogenetic estimation from DNA sequences with variable rates over sites: Approximate methods. J Mol Evol 39, 306–314 (1994).

49. Jombart, T. & Dray, S. adephylo: exploratory analyses for the phylogenetic comparative method. Bioinformatics 26, 1907–1909 (2010).

50. Pagel, M. Inferring the historical patterns of biological evolution. Nature 401, 877–884 (1999).

51. Orme, D. et al. caper: Comparative analyses of phylogenetics and evolution in R. Preprint at (2023).

