## Supplementary material for "Punctuational evolution is pervasive in distal site metastatic colonization": Figure S1.

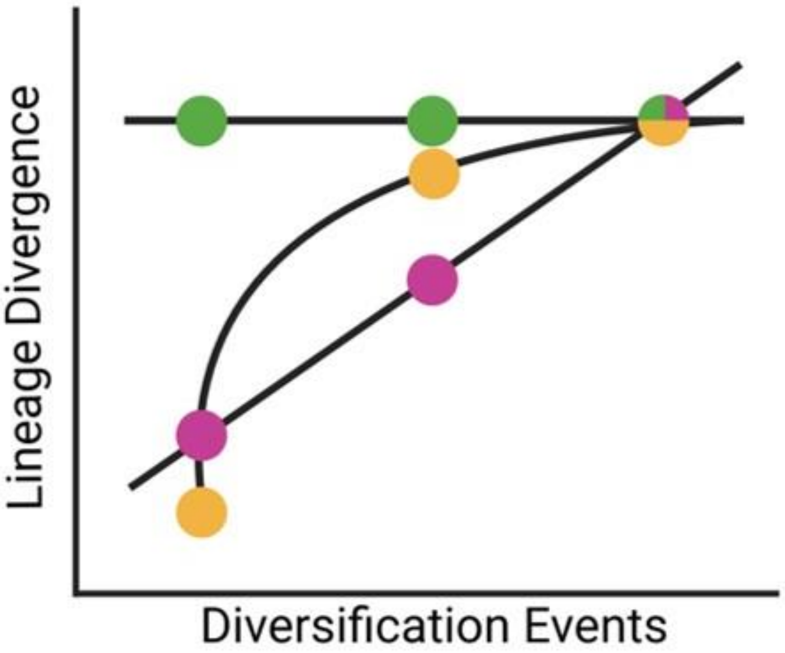


**Figure S1. Detecting the node density artifact.** The relationship between lineage divergence, the sum of the branch lengths from root-to-tip, and the number of lineage diversification events along the path, can be used to detect the node density artifact. A gradual model of evolution predicts no association between lineage divergence and diversification events (shown in green). In contrast, a punctuational model predicts a positive association between lineage divergence and diversification events (shown in pink). The node-density artifact is detected by fitting a curvilinear model (shown in yellow) and is deemed to present if the estimated exponent, $\delta$, is numerically greater than 1 (see Methods).
