## Supplementary material for "Punctuational evolution is pervasive in distal site metastatic colonization": Figure S2.

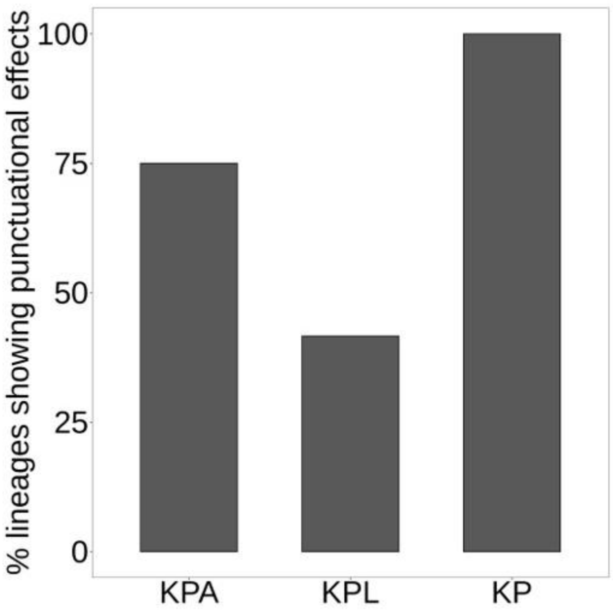


**Figure S2. Punctuational evolution is independent of the GEMM background.** The dataset consists of 18 lineages across 10 mice and three different genetically engineered mouse models (GEMMs): KP, KP, and KPL (see Methods). We found no association between the frequency of punctuational evolution and the specific mouse model ($p = 0.207$).
