## Supplementary material for "Punctuational evolution is pervasive in distal site metastatic colonization": Table S1.

| Lineage number | Metastatic | Number of cells | Node density artifact | % significant lineage diversification slope posterior | Punctuational effect | Mean punctuational contribution |
|---|---|---|---|---|---|---|
| 2 | Yes | 2421 | Yes | NA | NA | NA |
| 4 | Yes | 1612 | No | 100 | Yes | 0.011 |
| 5 | Yes | 1609 | No | 100 | Yes | 0.09 |
| 6 | Yes | 1213 | No | 97.5 | Yes | 0.072 |
| 7 | Yes | 1067 | No | 95.7 | Yes | 0.035 |
| 8 | Yes | 890 | Yes | NA | NA | NA |
| 9 | Yes | 849 | No | 100 | Yes | 0.04 |
| 10 | Yes | 468 | Yes | NA | NA | NA |
| 11 | Yes | 430 | No | 100 | Yes | 0.094 |
| 12 | Yes | 364 | No | 100 | Yes | 0.026 |
| 29 | No | 700 | Yes | NA | NA | NA |
| 35 | No | 603 | Yes | NA | NA | NA |
| 41 | No | 401 | No | 0 | No | NA |
| 42 | No | 348 | Yes | NA | NA | NA |
| 43 | No | 342 | No | 99 | Yes | 0.025 |
| 46 | No | 301 | No | 100 | Yes | 0.116 |
| 48 | No | 247 | No | 100 | Yes | 0.125 |
| 49 | No | 246 | No | 0.1 | No | NA |
| 52 | No | 213 | No | 1.3 | No | NA |
| 61 | No | 135 | No | 0 | No | NA |
| 65 | No | 118 | No | 0 | No | NA |
| 71 | No | 93 | Yes | NA | NA | NA |
| 72 | No | 90 | No | 10.9 | No | NA |
| 73 | No | 78 | No | 0 | No | NA |
| 74 | No | 44 | No | 0 | No | NA |

**Table S1. List of reconstructed single cell lineages and punctuational results.**
