## Supplementary material for "Punctuational evolution is pervasive in distal site metastatic colonization": Table S2.

| Lineage number | Number of cells | % significant lineage diversification slope posterior | Significant lineage diversification slope | % significant CNV slope posterior | Significant CNV slope |
|---|---|---|---|---|---|
| 4 | 1612 | 100 | Yes | 0 | No |
| 5 | 1609 | 100 | Yes | 3.4 | No |
| 6 | 1213 | 97.8 | Yes | 5.9 | No |
| 7 | 1067 | 95.7 | Yes | 0 | No |
| 9 | 849 | 100 | Yes | 0 | No |
| 11 | 430 | 100 | Yes | 0.5 | No |
| 12 | 364 | 100 | Yes | 0 | No |
| 41 | 401 | 0 | No | 0 | No |
| 43 | 342 | 98.9 | Yes | 0 | No |
| 46 | 301 | 100 | Yes | 0 | No |
| 48 | 247 | 100 | Yes | 0 | No |
| 49 | 246 | 0.1 | No | 0 | No |
| 52 | 213 | 1.7 | No | 0 | No |
| 61 | 135 | 0 | No | 0 | No |
| 65 | 118 | 0 | No | 0 | No |
| 72 | 90 | 12 | No | 0 | No |
| 73 | 78 | 0 | No | 0 | No |
| 74 | 44 | 0 | No | 0 | No |

**Table S3. Testing for an association between lineage divergence and copy number variations (CNVs)**
