## Supplementary material for "Punctuational evolution is pervasive in distal site metastatic colonization": Table S3.

| Lineage number | Number of cells | % significant lineage diversification slope posterior | Significant lineage diversification slope | % significant metastatic lineage diversification slope posterior | Significant metastatic lineage diversification slope |
|---|---|---|---|---|---|
| 4 | 1612 | 100 | Yes | 0 | No |
| 5 | 1609 | 100 | Yes | 1.8 | No |
| 6 | 1213 | 97.6 | Yes | 2.4 | No |
| 7 | 1067 | 95**.7** | Yes | 0 | No |
| 9 | 849 | 100 | Yes | 0 | No |
| 11 | 430 | 99.9 | Yes | 0 | No |
| 12 | 364 | 100 | Yes | 0 | No |

**Table S2. Testing for a metastatic specific association between lineage divergence and lineage diversification.**
