## Supplementary material for "Punctuational evolution is pervasive in distal site metastatic colonization": Table S4.

| Lineage number | Number of cells | Mean primary tumor punctuational contribution | Mean metastatic punctuational contribution |
|---|---|---|---|
| 4 | 1612 | 0.005 | 0.053 |
| 5 | 1609 | 0.021 | 0.101 |
| 6 | 1213 | 0.061 | 0.097 |
| 7 | 1067 | 0.013 | 0.162 |
| 9 | 849 | 0.026 | 0.089 |
| 11 | 430 | 0.057 | 0.064 |
| 12 | 364 | 0.028 | 0.011 |

**Table S4. The punctuational contribution within the primary tumor and distal site metastases for metastatic lineages.**
